# Modeling zero-inflated count data with glmmTMB

**DOI:** 10.1101/132753

**Authors:** Mollie E. Brooks, Kasper Kristensen, Koen J. van Benthem, Arni Magnusson, Casper W. Berg, Anders Nielsen, Hans J. Skaug, Martin Mächler, Benjamin M. Bolker

## Abstract

Ecological phenomena are often measured in the form of count data. These data can be analyzed using generalized linear mixed models (GLMMs) when observations are correlated in ways that require random effects. However, count data are often *zero-inflated*, containing more zeros than would be expected from the standard error distributions used in GLMMs, e.g., parasite counts may be exactly zero for hosts with effective immune defenses but vary according to a negative binomial distribution for non-resistant hosts.

We present a new R package, `glmmTMB`, that increases the range of models that can easily be fitted to count data using maximum likelihood estimation. The interface was developed to be familiar to users of the `lme4` R package, a common tool for fitting GLMMs. To maximize speed and flexibility, estimation is done using Template Model Builder (`TMB`), utilizing automatic differentiation to estimate model gradients and the Laplace approximation for handling random effects. We demonstrate glmmTMB and compare it to other available methods using two ecological case studies.

In general, glmmTMB is more flexible than other packages available for estimating zero-inflated models via maximum likelihood estimation and is faster than packages that use Markov chain Monte Carlo sampling for estimation; it is also more flexible for zero-inflated modelling than `INLA`, but speed comparisons vary with model and data structure. Our package can be used to fit GLMs and GLMMs with or without zero-inflation as well as hurdle models. By allowing ecologists to quickly estimate a wide variety of models using a single package, glmmTMB makes it easier to find appropriate models and test hypotheses to describe ecological processes.

## 1. Introduction

Ecological phenomena are often measured in the form of discrete count data, e.g., the number of times that owl nestlings beg for food (Roulin & Bersier, 2007), counts of salamanders in streams (Price et al., 2016), or counts of parasite eggs in fecal samples of sheep (Brown et al., 2012). These counts are often analyzed using generalized linear models (GLMs) and their extensions (O’Hara & Kotze, 2010; Wilson & Grenfell, 1997). GLMs quantify how expected counts change as a function of predictor variables, e.g., nestlings change their behavior depending on which parent they interact with (Roulin & Bersier, 2007), salamander abundance decreases in streams affected by coal mining (Price et al., 2016), and helminth infection intensity in sheep varies with age and genotype (Brown et al., 2012). Repeated measurements on the same individual, the same location, or observations taken at the same point in time are often correlated; this correlation can be accounted for using random effects in generalized linear mixed models (GLMMs; Bolker et al., 2009; Bolker, 2015).

These types of count data are commonly modeled with GLMs and GLMMs using either Poisson or negative binomial distributions. For the Poisson distribution, the variance is equal to the mean. When data are overdispersed — meaning the variance is larger than the mean — they are often instead modeled using the negative binomial distribution, which can be defined as a mixture of Poisson distributions with Gamma-distributed rates. For the Poisson and negative binomial distributions, the expected number of zeros decreases as the mean increases. However, when multiple processes underlie the observed counts — which is almost ubiquitous in biology — the counts can contain many zeros even if the mean is much greater than zero. For example, observed counts of salamanders could be zero if a stream is uninhabitable due to mining waste, or the count could be any integer from zero to infinity depending on other qualities of the stream affecting the population density and the salamanders’ ability to hide from researchers (Price et al., 2016). Zero-inflated GLMs allow us to model count data using a mixture of a Poisson or negative binomial distribution and a structural zero component, i.e., extra zeros. Models that ignore zero-inflation, or attempt to handle it in the same way as simple overdispersion, yield biased parameter estimates (Harrison, 2014).

Many biologists use the statistical computing environment R and its contributed packages to organize, model, and graph their data (R Core Development Team, 2016). In R, there are five main packages available for modeling zero-inflated data: pscl, `INLA, MCMCglmm, glmmADMB`, and brms (Table 1; Zeileis et al., 2008; Rue et al., 2009; Hadfield, 2010; Skaug et al., 2012; Bürkner, in press). The `pscl` package can fit zero-inflated GLMs with predictor variables on the zero-inflation using maximum likelihood estimation (Zeileis et al., 2008). For example, `pscl` can be used to test the hypothesis that sheep fecal egg counts depend on age and extra zeros depend on genotype. However, `pscl` cannot model the correlation within individuals if they are sampled repeatedly; this phenomenon requires random effects. Omitting random effects and thereby ignoring correlation makes statistical tests anti-conservative (Bolker et al., 2009; Bolker, 2015). The glmmADMB package can fit zero-inflated GLMMs that contain random effects to account for correlation among observations (Skaug et al., 2012). However, it cannot fit models with predictor variables in the zero-inflation part of the model; thus, it is only appropriate for limited cases where all observational units (e.g., individual sheep) have an equal probability of producing a structural zero. `INLA` has the same limitation as glmmADMB. The MCMCglmm and brms packages can fit zero-inflated GLMMs with predictors of zero-inflation, but they are relatively slow because they require Markov chain Monte Carlo (MCMC) sampling (Bürkner, in press; Hadfield, 2010).

**Table 1.** Features implemented in glmmTMB and other packages that are used for modeling zero-inflated count data. `lme4` and `mgcv` are not included because they can only estimate zero-inflation when wrapped in an iterative algorithm (Minami et al., 2007; Bolker et al., 2013).

| Feature | glmmTMB | glmmADMB | pscl | MCMCglmm | brms | INLA |
| --- | --- | --- | --- | --- | --- | --- |
| predictors of zero-inflation | ✓ |  | ✓ | ✓ | ✓ |  |
| predictors of dispersion | ✓ |  |  |  |  |  |
| zero-truncated distributions | ✓ | ✓ | ✓ | ✓ | ✓ | ✓ |
| `nbinom2` distribution | ✓ | ✓ | ✓ |  | ✓ | ✓ |
| `nbinom1` distribution | ✓ | ✓ |  |  |  |  |
| weights | <sup>1</sup> | ✓ | ✓ |  | ✓ | ✓ |
| offsets | ✓ | ✓ | ✓ | <sup>2</sup> | ✓ | ✓ |
| random effects (RE) | ✓ | ✓ |  | ✓ | ✓ | ✓ |
| various RE structures | ✓<sup>3</sup> |  |  | ✓ | ✓ | ✓ |
| maximum likelihood estimation | ✓ | ✓ | ✓ |  |  |  |
| MCMC sample a fitted model | ✓<sup>4</sup> | ✓ |  | ✓ | ✓ | ✓ |
| multivariate responses |  |  |  | ✓ | ✓ |  |

Notes: <sup>1</sup> Weights are often used to reduce the influence of some observations over others, e.g., (Gurevitch & Hedges, 1999); `glmmTMB`'s dispersion formula can be used to model heteroskedasticity, but might not fill other roles of weights. This feature may be added in future versions. <sup>2</sup> Offsets can be implemented using priors. <sup>3</sup> See `vignette("covstruct")` for details. <sup>4</sup> See `vignette("mcmc")` for details.

Here we present a new R package, glmmTMB, that estimates GLMs, GLMMs and extensions of GLMMs including zero-inflated GLMMs using maximum likelihood. The ability to fit these types of models quickly and using a single package will make it easier for biologists to find the best model to explain patterns in their data. We demonstrate the package using two examples. We use an example of salamander abundance to show how to fit and compare zero-inflated and hurdle GLMMs and then how to extract results from a model. We use a classic example of owl nestling behavior to compare the timing and parameter estimates from glmmTMB to other R packages.

## 2. Implementation of glmmTMB

The design goal of glmmTMB is to extend the flexibility of GLMMs in R while maintaining a familiar interface. To maximize flexibility and speed, `glmmTMB`’s estimation is done using the `TMB` package (Kristensen et al., 2016), but users need not be familiar with `TMB`. We based `glmmTMB`’s interface (e.g., formula syntax) on the `lme4` package — one of the most widely used R packages for fitting GLMMs (Bates et al., 2015). Like `lme4`, `glmmTMB` uses maximum likelihood estimation and the Laplace approximation to integrate over random effects; unlike `lme4`, `glmmTMB` does not have the alternative options of doing restricted maximum likelihood (REML) estimation nor using Gauss-Hermite quadrature to integrate over random effects (Bolker et al., 2009; Bolker, 2015). REML may be added to `glmmTMB` in the future. The underlying implementation using `TMB` is the major divergence from `lme4` and provides `glmmTMB` with a speed advantage when estimating non-Gaussian models (Appendix A) and greater flexibility in the types of models it can fit (Table 1). The flexibility of `glmmTMB` enables users to fit and compare many varieties of models with assurance that the loglikelihood values are calculated in comparable ways. Comparing likelihoods of models fit by multiple packages is not advisable because some packages drop constants from the log-likelihood calculations while others do not.

A `glmmTMB` model has four main components: a conditional model formula, a distribution for the conditional model, a dispersion model formula, and a zero-inflation model formula. Simple GLMs and GLMMs can be fit using the conditional model while leaving the zero-inflation and dispersion formulas at their default values. The mean of the conditional model is specified using a two-sided formula with the response variable on the left and predictors on the right, potentially including random effects and offsets. This formula uses the same syntax as `lme4`. For example, if salamander counts vary by species (spp) and vary randomly by site, then the formula for the dependence of mean count on species could be count*∼*spp + (1|site).

The distribution around the mean of the conditional model is specified using the argument family. For count data, the distribution will typically be either Poisson, negative binomial, or binomial, but we do not discuss details of the binomial here, because modeling zero-inflation is more common with Poisson and negative binomial distributions. The Poisson and negative binomial distributions use a log link by default, but identity links can be specified as in `family=poisson(link=“identity”)`. `glmmTMB` provides two parameterizations of the negative binomial which differ in the dependence of the variance (*σ*^2^) on the mean (*μ*). For family=nbinom1, the variance increases linearly with the mean as *σ*^2^ = *μ*(1 + *α*), with *α >* 0; for family=nbinom2, the variance increases quadratically with the mean as *σ*^2^ = *μ*(1 + *μ/θ*), with *θ >* 0 (Hardin & Hilbe, 2007).

With the default dispersion model (dispformula=*∼*1), the dispersion parameter (e.g., *α* or *θ* for the negative binomial distribution) is identical for each observation. Alternatively, the dispersion parameter can vary with fixed effects; in this case, the dispersion model uses a log link. The dispersion model can be used to account for heteroskedasticity. For example, if the response is more variable (relative to the mean) as the year progresses, then a model with either negative binomial distribution might use the one-sided formula dispformula=*∼*DOY where DOY is the day of the year. A description of the dispersion parameter for each distribution can be accessed by typing `?sigma.glmmTMB` in R. The Poisson and binomial distributions do not use a dispersion parameter.

The zero-inflation model describes the probability of observing an extra (i.e., structural) zero that is not generated by the conditional model. Zero-inflation creates an extra point mass of zeros in the response distribution; the overall distribution is a mixture of the conditional model and zero-inflation model (Lambert, 1992; Rhodes, 2015). The zero-inflation probability is bounded between zero and one by using a logit link on the zero-inflation model. For example, if salamanders were expected to emerge on some date after observations began being collected, then the model could include the one-sided formula ziformula=*∼*DOY. The probability of producing a structural zero can be modeled as equal for all observations with ziformula=*∼*1.

## 3. Installation

The package is available from The Comprehensive R Archive Network (CRAN) via the command `install.packages(“glmmTMB`”). Development versions are available from GitHub and can be installed using devtools (Wickham & Chang, 2016). Current details for installing development versions should be accessed on the GitHub page https://github.com/glmmTMB/glmmTMB.

## 4 Case studies

To illustrate how to use `glmmTMB` and to compare it to other packages, we applied it to two publicly available ecological data sets, both of which are included in the package.

### 4.1. ABUNDANCE OF SALAMANDERS IN STREAMS

We analyzed counts of salamanders in streams to demonstrate how to use features of `glmmTMB` to do model selection and to output model results. Salamanders of seven different species and life stages were repeatedly sampled and replaced four times at 23 sites in streams (Fig B.1). Some of the streams were impacted by mountaintop removal and valley filling from coal mining. The data set contains covariates that may affect the habitat suitability of a site and the ability of researchers to capture salamanders that inhabit the site (Price et al., 2016, 2015). Price et al. analyzed the data using a Bayesian model with an ecological and a sampling component; abundance was estimated with a hurdle Poisson model and then observations were modeled as binomial samples from the abundance. We fit GLMMs, zero-inflated GLMMs, and hurdle models to this data with Poisson and negative binomial distributions on the conditional model. For simplicity, we did not consider all variables that may affect the counts. As a null model, we assumed that counts varied by species (spp) and randomly by site (site). We fit models where the mean count additionally depended on mining (mined). Our full zero-inflated GLMMs allowed both the conditional and zero-inflation models to differ between mined and unmined sites. The full zero-inflated negative binomial GLMM was fit using the command `glmmTMB`(count*∼*spp * mined + (1|site), ziformula=*∼*spp * mined, family=nbinom2, data=Salamanders). As is generally the case for model formulas in R, the * indicates an interaction plus main effects. We used Akaike information criteria to compare all models. Then we illustrated the assessment of a zero-inflated model using the summary, predict, and simulate functions (Figs B.2, B.3, B.4, and B.5). See Appendix B for details, including code. Then, we measured the time required to fit the most parsimonious model across multiple packages. We performed three sets of timing benchmarks: (1) on simulated data with the same structure as the original data, (2) on the original data replicated to create more observations per site and the same number of sites, (3) on simulated data with increasing number of sites and the same number of observations per site. Benchmarks were run in parallel using parLapply (R Core Development Team, 2016) on a MacBook with 2.7 GHz Intel Core i7 processor, 8 cores, and 16 GB memory.

Of the models we considered, the most parsimonious was a negative binomial GLMM that allowed salamander counts to vary with species, mining, and their interaction. These results agree with earlier work showing that the negative binomial distribution is often the best fit for abundance data (Warton, 2005). Fitting this model to simulated data with the same structure as the original data was, on average, equally fast in `glmmTMB` and `INLA`, 26 times slower with glmmADMB, 30 times slower with `lme4`, and 274 times slower with brms (Fig A.1). Fitting the model to the original data replicated to have more observations per site required, on average, half the time to fit with `INLA` compared to `glmmTMB`, 22 times as long with glmmADMB, 30 times as long with `lme4`, and 59 times as long with brms (Fig A.2). For simulated data sets with more sites (i.e., more random effect levels), `INLA`’s estimation time accelerated such that it was faster than `glmmTMB` for data sets with fewer than 5888 sites, but slower for those with more sites (Fig A.3). Benchmarking nuances could be investigated in more detail in future studies.

Two zero-inflated negative binomial models did not converge; they both allowed zero-inflation to vary by species but not with mining, which could be unreasonable. General model convergence issues are discussed in `vignette(“troubleshooting”)`.

### 4.2. BEGGING BEHAVIOR OF OWL NESTLINGS

To further compare R packages for fitting zero-inflated GLMMs, we analyzed counts of begging behavior by owl nestlings. The full analyses can be found in Appendix C. This example previously appeared in Zuur et al. (2009) and Bolker et al. (2013) and was originally published by Roulin & Bersier (2007). We compared the estimates of fixed effects and the amount of time required for fitting the same model in `glmmTMB`, glmmADMB, `INLA`, brms, and MCMCglmm (Skaug et al., 2012; Rue et al., 2009; Bürkner, in press; Hadfield, 2010). For brms and MCMCglmm, we used the default number of iterations, burn-in samples, and thinning. In each package, we fit zero-inflated Poisson models with five fixed effects, an offset term, and one random effect. We allowed zero-inflation to vary with food treatment and vary randomly with nest. See Appendix C for details of these methods, including code.

On a standard laptop computer, it took approximately five seconds to fit a zero-inflated Poisson GLMM in `glmmTMB`; MCMCglmm took four times as long as `glmmTMB`, followed by glmmADMB (eight times as long) and brms (27 or 14 times as long, depending on precompilation). `INLA` was slightly faster than `glmmTMB`. Estimates and confidence (or credible) intervals (CI) from brms and `INLA` were nearly identical to those of `glmmTMB`, when running the Bayesian models with flat priors (Figs C.1 and C.2). Estimates from glmmADMB and `glmmTMB` differed slightly, because glmmADMB uses numerical adjustments to increase the probability that a model will converge and these change the objective function by a small amount (Fig C.1). Estimates and CI from MCMCglmm differed from those of `glmmTMB` because the Bayesian model included informative priors (Figs C.1 and C.2); some priors were necessary to force part of the model to be treated as an offset rather than as a standard predictor variable.

## 5. Conclusions

We have introduced an R package that can quickly estimate a variety of models including GLMs, GLMMs, zero-inflated GLMMs, and hurdle models. By providing this flexibility in a single package, we reduce the need for researchers to learn multiple packages. Another benefit is that models estimated with a single package can be compared using likelihood-based methods including information criteria.

## Acknowledgements

Thanks to S. Price for providing the data on salamanders and for helpful comments on the manuscript. Thanks to A. Zuur for providing the data on owl nestlings. This work was supported by grants from the Swiss National Science Foundation (#IZ320Z0 161670) and from the AD Model Builder Foundation to MEB.

